# Serotonergic neurons in the dorsal raphe regulate visual attention

**DOI:** 10.1101/2024.09.29.615662

**Authors:** Jonas Lehnert, Kuwook Cha, Julia Forestell, Kerry Yang, Xinyue Ma, Jonathan Britt, Anmar Khadra, Erik P. Cook, Arjun Krishnaswamy

## Abstract

Visual attention enhances the neural representation of salient stimuli within the visual cortex. It is generally thought that this enhancement is driven by glutamatergic feedback from frontal cortical areas. Here we report the unexpected observation that dorsal raphe (DR) derived serotonin (5HT) controls visual attention. We developed a behavioral model that captured the way mice allocated attention to cued and uncued visual locations and features. Simultaneous photometry showed reduced DR activity when mice deployed attention to the cued locations and features, whereas high DR activity was observed when mice were less attentive. Optogenetic excitation of DR-5HT neurons impaired attention to the cue and degraded behavioral performance, while optogenetic suppression improved attention and performance. A genetically encoded sensor of 5HT release showed reduced 5HT levels in visual cortex when mice attend and detect stimuli. These results demonstrate that DR-5HT neurons are members of the brain’s attentional circuit and suggest that 5HT is a novel biological carrier of visual attention.

## Introduction

Visual attention enhances neural representations of visual locations and features that are the most relevant to the task at hand^1–3^. Neural circuits in visual cortex (V1) mediate this enhancement but the way they are driven according to internal needs for visual attention is not clear^1,4^. Some attention control signals feedback to V1 from frontal cortical areas^1,4–11^, however, cortically-released neuromodulators are also believed to play a major uncharacterized role^12–18^. Of these factors, serotonin (5HT) is perhaps the most intriguing because cortical 5HT release is influenced by frontal areas^19–22^, implicated in internal states, and strongly modulates visual responses^23–29^. Yet we know little about the contribution of 5HT to visual attention.

Cortical 5HT comes from projection neurons residing in the dorsal raphe (DR) that encode states, such as reward^30,31^ or patience^32–34^. DR neurons innervate many cortical areas including visual and frontal areas^19–22^. Recent optogenetic studies revealed powerful effects of DR-5HT on mouse visual cortical neurons. In one study, DR stimulation suppressed both spontaneous and visually evoked activity of V1 measured with wide-field imaging^24^. Stimulating DR-5HT neurons in a second study suppressed responses of V1 neurons to their preferred stimuli^23^. These results echo older work in non-human primates showing that 5HT iontophoresis decreased response gain of V1 neurons^25,26,35^. Collectively, these studies show that 5HT modulates visual responsiveness, but is there a link between DR-5HT and visual attention?

To address this question, we optically monitored and manipulated DR-5HT signals in mice performing a cued detection task and analyzed their attention to visual features and locations. Using fiber photometry, we demonstrate that DR neurons encode various task events/outcomes and that their activity drops when mice selectively attend to cued locations. Low DR activity predicted greater visual attention and improved performance whereas high DR activity was associated with less attention and poorer performance in our task. Optogenetic excitation of DR neurons prior to stimulus presentation deteriorated attention and performance whereas optogenetic suppression improved attention and performance. Using a genetically encoded 5HT sensor, we show that low 5HT in V1 was correlated with enhanced attention and detection performance. These results define DR 5HT neurons as a new regulator of visual attention.

## Results

We recently developed a cued detection task that controls contextual signals related to attention, but also allow unbiased measure of other internal contexts that change perceptual sensitivity and behavioral performance. Briefly, water restricted mice were head-fixed atop a platform facing a pair of angled screens and tasked to search for 3 white vertical bars that emerge from dynamic checkerboard noise (**Fig. 1a, top**). We varied trial difficulty by varying the amount of noise combined with the 3-bars (**Fig. 1a, bottom**). Trials began with a tone and static checkerboard on one monitor which acted as an audio-visual cue for the eventual location of the 3-bar grating (**Fig. 1b**). Mice initiated trials by licking a spout, which caused the cue to fade while zero-coherence dynamic checkerboard noise appeared on both screens for a random period (**Fig. 1b**, *delay period*) until the 3-bar grating appeared; the probability of grating appearance was a constant flat hazard function. Because gratings were noisy and appeared

**Figure 1.**
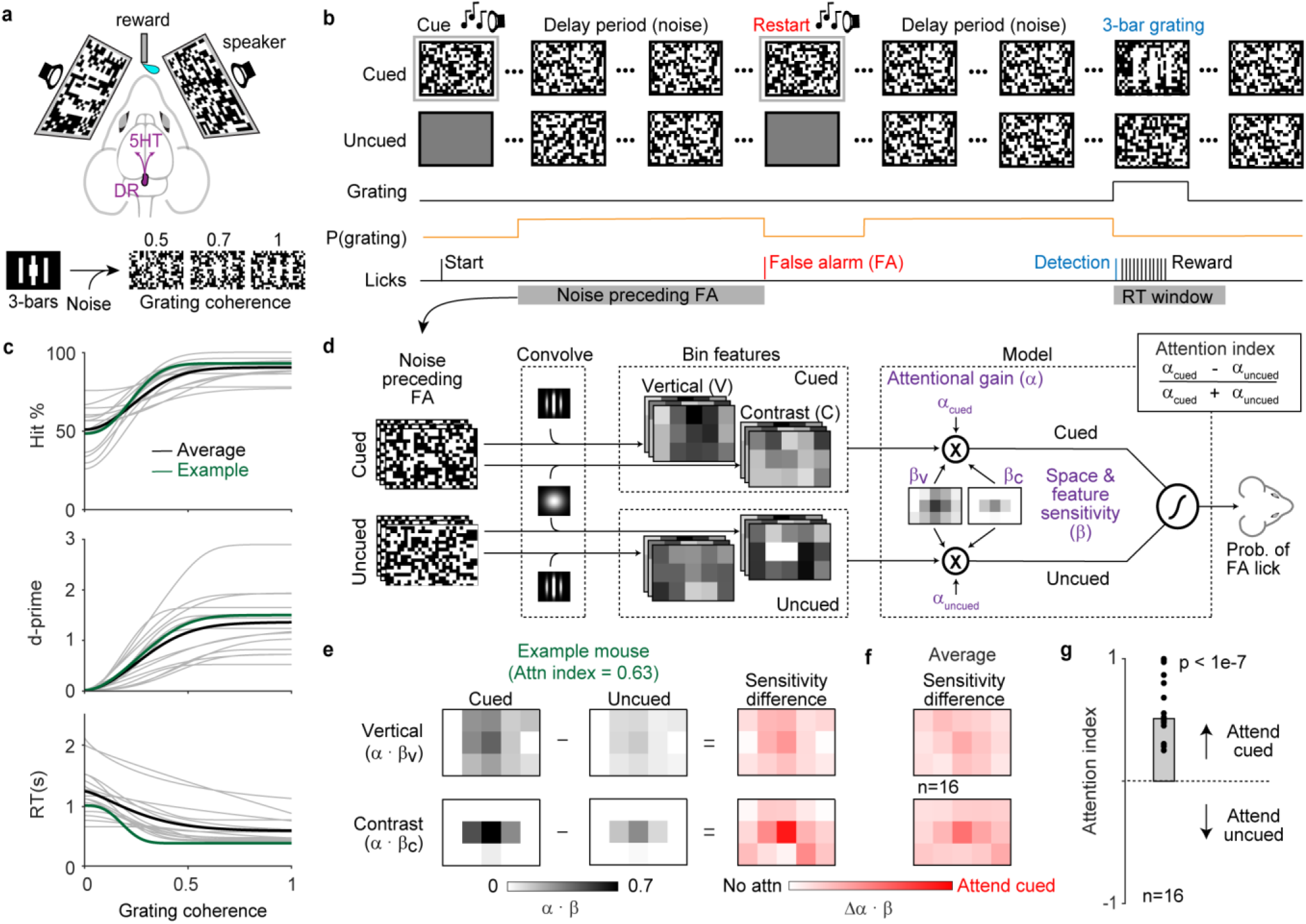
Task to measure attention induced changes in visually guided behavior. **a**, Top: cartoon illustrating DR 5HT neuromodulation in our visually cued task. Head-fixed mice faced angled visual displays (100×64^°^ of visual angle) and licked a spout when they detected 3-bar gratings to obtain a reward. Bottom: example 3-bar gratings shown to mice produced by combining 3-bars with different amounts of checkerboard noise. Coherence = 1, all 3-bar checkers are white. Coherence = 0, all 3-bar checkers are noise. **b**, Trials began with a static checkerboard and audiovisual cue that indicated the eventual location of the 3-bar grating. The cue quickly faded after mice licked (Start) and was replaced by dynamic checker-board noise (30 Hz) presented for a randomly chosen delay period. A lick within a 1.4 sec reaction time (RT) window that followed the 3-bar grating (Detection) resulted in a reward (Reward). Cue and coherent stimulus switched screens approximately every 25 trials. A lick during the delay period (False Alarm, FA) restarted the cue and trial. Such FA restarts recurred until the delay period was lick free. **c**, Psychometric curves from trained mice relating %hits, d’, and RT to stimulus coherence. Gray lines are individuals, black is the mean (∼3000 trials/mouse; n=13) and green is an example mouse. **d**, Reverse correlation model to estimate behavioral attention to visual features and locations. Individual checkerboard frames are (1) convolved with vertical Gabor filters or 2D Gaussians to obtain vertical and contrast energy maps; (2) binned across 15 locations; and (3) weighed depending on whether energies come from cued or uncued sides; (4) A logistic regression model fit parameters for behavioral sensitivity to the binned vertical and contrast energy maps (*B*_V_ and *B*_C_) and attentional gain (α_Cued_ and α_Uncued_) to predict the probability of a FA lick. **e**, Model results from example mouse (green in Panel c). Behavioral attention is reported as either an attention index based on attentional gain, or differences in the scaled sensitivities to cued and uncued sides. Color scale = 0 - 0.7. **f**, Population sensitivity difference maps show enhanced attention to the center of the cued screen before FA licks. Color scale = 0 - 1 **g**, Population attention indices indicate attention was directed to the cued screen before FA licks (one sample t-test) unpredictably, our design forced mice to continually evaluate the checkerboard noise for evidence of the 3-bar grating. Licks to grating onset (*hits*) led to a liquid reward (**Fig. 1b-c**). Licks to a zero-coherence grating were unrewarded and used to compute psychometric measures. Mice showed reliable detection (d-prime ≥ 1) of very weak gratings (coherence = 0.3) after ∼3 weeks of training (**Fig. 1c**).

False-alarm licks (FAs) during the delay period caused trials to restart until the wait period was lick-free (**Fig. 1b**). We previously used such FAs to reverse correlate the sensitivity of mice to visual locations and features, and found that they attended a 30degree patch of the cued monitor for vertical patterns and local contrast^36^. Here, we simplified this model to reflect these prior observations. Briefly, our model convolved checkerboard patterns with a vertical Gabor to extract vertical energy or a 2D Gaussian filter to extract local contrast energy (**Fig. 1d**, *convolve*). Next, we binned both kinds of energy map in a 5×3 grid encompassing the entire checkerboard (*bin features*, bin = ∼10^°^). Finally, a logistic regression model weighed the binned energies on the cued and uncued screens to predict FA licks (*model*). Fitting the model to each mouse provided two sets of parameters: 1) Weights that captured attentional gain to the cued versus the uncued screen (α_Cued_ and α_Uncued_); 2) Weights that captured sensitivity to both vertical energy (β_V_) and contrast energy (β_C_) across space.

This model had significant predictive power when fit to each mouse (**Extended data Fig. 1a**) and quantified attention to cued versus uncued screens as either a difference in sensitivity (**Fig. 1e, f**) or as an attentional-gain index (**Fig. 1e, g**). The model’s sensitivity difference (Δα ⋅ β_V_ and Δα ⋅ β_C_) shows that mice attended to stimulus energies near the center of the cued screen (**Fig. 1f**), consistent with our prior observation of an attentional hotspot in this location^36^ (**Extended Data Fig. 1b**). The attentional index showed mice consistently weighed cued stimulus energies ∼60% more than those of the uncued side (**Fig. 1g)**. Taken together, these behavioral measures show that mice followed the cue to attend and detect the noisy 3-bar grating.

### DR neurons encode several events and outcomes of our visually guided task

Next, we used fiber photometry to investigate how the DR reacted to events and outcomes in our task (**Fig. 2a**). Briefly, we crossed Cre-lines that mark all DR-5HT neurons with lines containing a Cre-dependent transgene for the genetically encoded calcium indicator GCaMP6f. Most experiments were conducted with *epet1-Cre*, but a few initial studies used Cadherin13-CreER (*Cdh13-CreER*). Sequencing atlases predict that both genes label similar subsets of DR-5HT neurons^19–21,37^, and consistent with this, both lines showed a similar percent of GCaMP6f+ DR neurons expressing the 5HT synthesis enzyme, tryptophan hydroxylase 2 (Tph2, **Fig. 2b-e, Extended Data Fig. 2a-c**). Both lines also produced similar task-related signals, so we grouped them together (**Extended Data Fig. 2d-g**).

**Figure 2.**
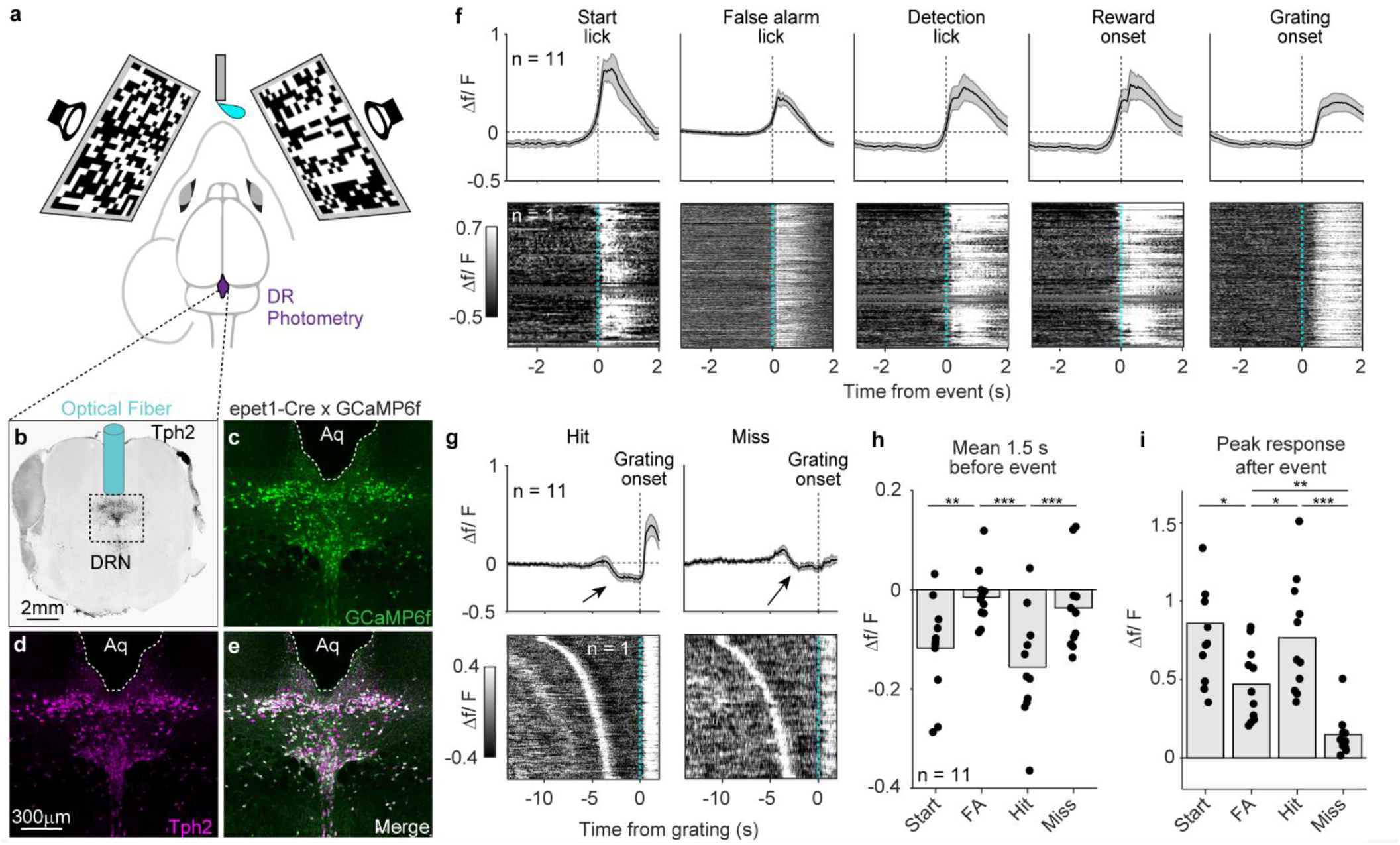
DR neural signals encode trial events and outcomes. **a**, Cartoon shows fiber photometry from DR while mice detect 3-bar gratings. **b**, Coronal brain section immunostained with antibodies against tryptophan hydroxylase 2 (Tph2, black); optical fiber position indicated. **c-e**, Coronal DR section from an epet1-Cre x GCaMP6f mouse immunostained for GCaMP6f (c), Tph2 (d), and a merge (e). **f**, Mean GCaMP6f signal (top) and an example mouse raster of raw recordings (bottom) from DR-5HT neurons. Traces are aligned to the indicated trial events (n=11). **g**, Mean photometry signal and example raster of raw responses aligned to grating onset for hit and miss trial outcomes. DR is suppressed before grating presentation on hit trials (arrows). Rasters are sorted based on the length of the delay period before the grating onset. **h-i**, Mean GCaMP signal before the indicated events (h) and average response amplitude of these events (i). * = P <0.05; ** = P < 0.01; *** = P < 0.001, paired t-test.

Task-evoked events drove positive signals from DR-5HT neurons that varied in their kinetics and magnitude depending on their association with start, false alarm, detection, or reward licks (**Fig. 2f, h-i**). Grating presentation did not immediately produce a positive signal, consistent with the idea that DR does not receive strong visual input (**Fig. 2f**, *grating onset*). We also saw a small peak aligned to reward delivery following detection lick, consistent with prior work associating DR with reward^30,38^ (**Fig. 2f** *detection lick*). Changes in fluorescence unrelated to DR neural activity were controlled by recording calcium-sensitive and -insensitive GCaMP fluorescence by simultaneously imaging with 470nm and 405nm excitation wavelengths (*see methods*).

DR signals during hit and miss trials differed in a few ways (**Fig. 2g-i**). Hits were associated with positive responses after grating presentation whereas misses were not (**Fig. 2g**). We noticed that hits were often preceded by low DR activity (arrow) that appeared following licks to the cue and persisted until grating onset (**Fig. 2g**, *hit*). In contrast, misses were preceded by higher DR activity (**Fig. 2g**, *miss*).

This anticipatory DR low signal differs from prior work on task-related DR signals in two ways. First, it is an order of magnitude faster than previous reports showing minute-long DR high and low dynamics^31^; and second, it covaried with hits and misses (**Fig. 2g-i**). Our analysis of slow DR dynamics confirmed this view and showed no correlation between minute-long DR signals and task performance (**Extended data Fig. 3**). Thus, DR-5HT neurons encoded several events/outcomes of our visually guided task and showed an unexpected transient low signal that preceded grating detection.

### DR neural activity correlates with detection performance and attention

The lower DR activity during the delay period before hits (**Fig. 2g**) is also when mice deployed attention to the cued screen (**Fig.1f-g**). Given this and prior work that showed that 5HT modulates visual responses^24–27,35^, we next asked how DR activity is linked to behavioral performance.

We first separated trials into low and high groups based on the average DR activity before grating presentation (**Fig. 3a**). Next, we computed behavioral performance separately from DR-5HT low and high trials. Psychometric measures (% hits, d-prime, threshold) from DR low trials showed enhanced detection relative to DR high trials (**Fig. 3b-d**); no changes were seen in reaction time. These differences arose without changes in response bias, lapse rate, or criterion (**Fig. 3e**), indicating that DR activity predicts grating sensitivity rather than impulsivity, engagement, or strategy. The enhanced detection during low DR activity was most pronounced for trials where the stimulus coherence was low (**Fig. 3c-d**). This is consistent with work showing the strongest effects of attention are usually observed when detecting weak stimuli^39^.

**Figure 3.**
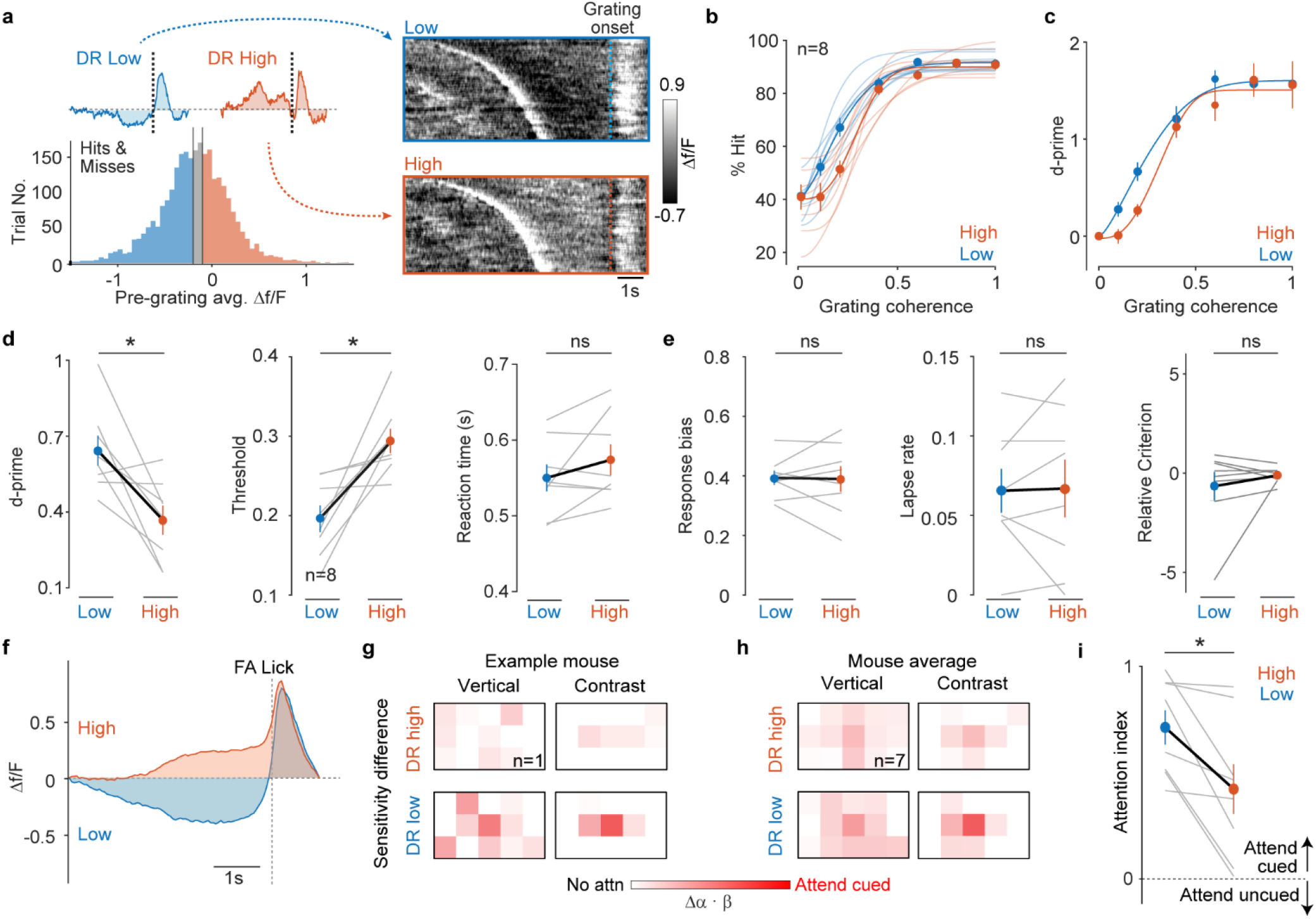
DR neural activity predicts attention and detection of the 3-bar grating. **a**. Histogram of mean DR signal preceding grating presentation divided into low- and high-activity groups for one mouse. Insets shows mean response, and rasters show raw responses of low and high groups aligned to grating onset. **b-c**, Psychometric curves of % hits (b) and d-prime (c) versus stimulus coherence for DR high and low groups. Mice perform with higher sensitivity in DR-low trials than in DR-high trials. Light lines are individual mice, thick lines are the mean. **d**, Mean d-prime at 0.2 coherence, psychometric threshold, and reaction time (RT) for DR high and low groups. **e**, Response bias, lapse rate, and criterion for DR high and low groups. Threshold is significantly reduced and d-prime significantly elevated in DR-low trials. **f**, Average DR high and low signals 1.5 seconds before a false alarm lick. **g-h**, Sensitivity difference maps for DR high and low in an example mouse (g) and averaged across 7 mice (h). DR low enhances attention to cued features and locations (n =7). Example mouse color scales: DR high = 0 - 0.2; DR low = 0 - 1. Mouse average color scales are 0 – 1 for DR high and low. **i**, Average attentional index for DR high and low states. Attention is significantly increased when the DR activity is low. * = P < 0.05, Wilcoxon signed-rank test.

Given this, we used our attentional gain model (**Fig. 1d**) to analyze FA licks preceded by either high or low DR activity (**Fig. 3f**). Sensitivity difference maps during DR low trials were much stronger than those computed from DR high. They also showed enhanced sensitivity to vertical and contrast energies at the center of the cued screen (**Fig. 3g-h**). Moreover, average attentional gain was ∼25% larger when the DR was low versus when it was high (**Fig. 3i**). Together, these results show that DR-5HT activity is inversely correlated with attention to visual locations and features that provide evidence of the 3-bar grating.

### Optogenetic manipulation of DR neurons changes animal behavioral performance and attention

The correlation between attentional gain and DR-5HT activity prior to detection strongly suggested that DR-5HT regulates attention. To test this hypothesis, we selectively expressed excitatory and inhibitory optogenetic actuators in DR-5HT neurons (**Fig 4a-b**) and asked whether suppressing or elevating their activity could alter attention and detection behavior (**Fig. 4c**). Briefly, mice expressing light-activated inhibitory (Jaws) or excitatory (ChR2) opsins in DR-5HT neurons were implanted with optical fibers and fully trained on the task before optogenetic manipulations began. Three seconds of light from a LED was used to drive either Jaws or ChR2 beginning two seconds before grating presentation (**Fig. 4d**). If mice produced false-alarm licks during the stimulation window, the LED was stopped, the cue was shown and the trial restarted (**Fig. 4d**, *Restart*). Thus, our optogenetic data naturally contained periods before FA licks with either the LED on or off. This allowed us to compare attention across three conditions: FAs in trials prior to stimulation (Ctrl), FAs directly preceded by stimulation (LED on), and FAs that were not preceded by stimulation (LED off).

**Figure 4.**
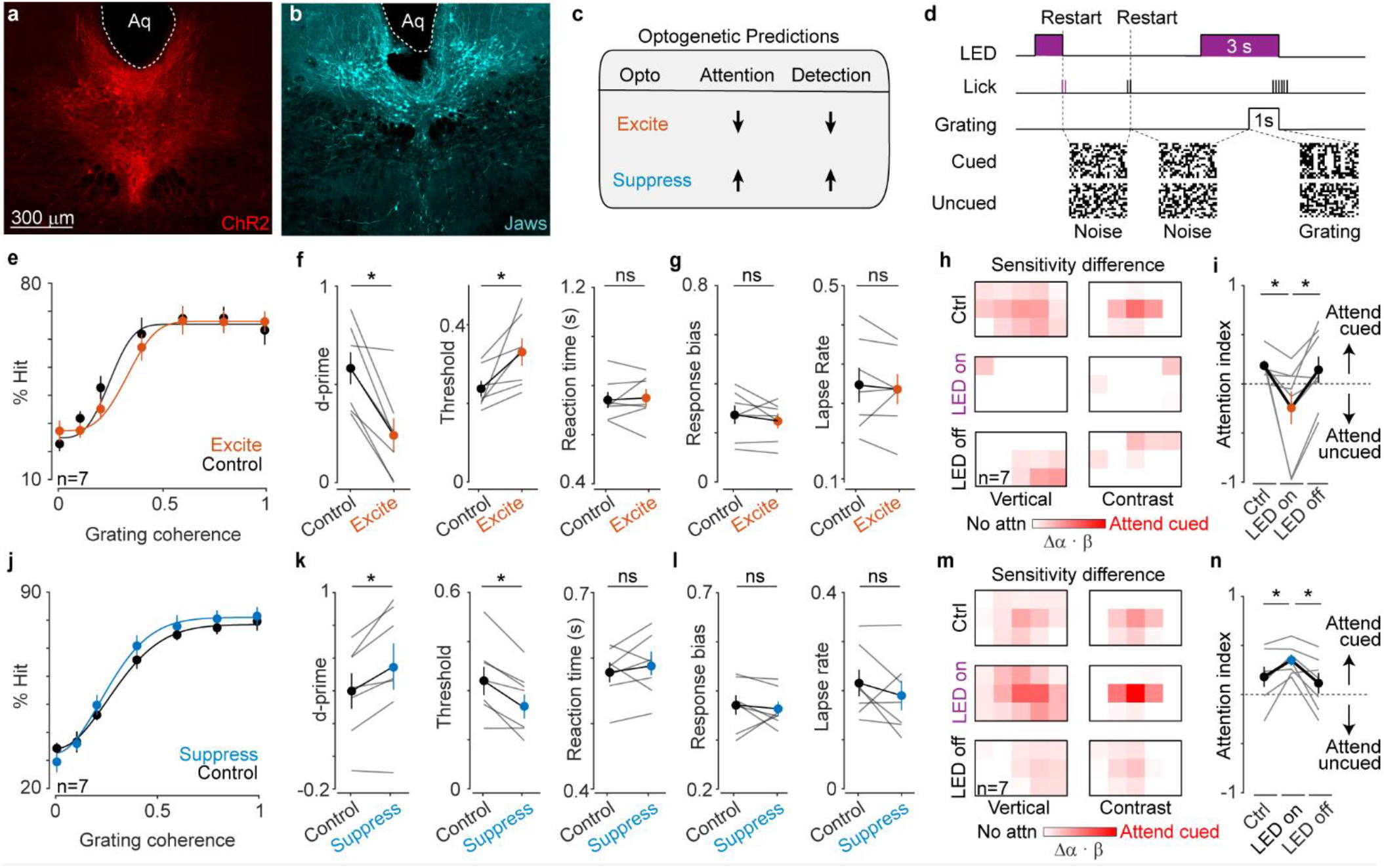
DR state regulates attention to visual features and space. **a-b**. Image of DR neurons expressing channelrhodopsin (a, ChR2) and Jaws (b). **c**. Optogenetic excitation (ChR2) and inhibition (Jaws) should suppress or enhance attention and detection of the 3-bar grating, respectively. **d**. A 20 Hz train of light pulses began 2 seconds prior to grating presentation and stopped when the grating disappeared. False alarms occurring during LED stimulation cause the trial to restart. **e**. % Hits versus stimulus coherence for unstimulated (Control) and ChR2-evoked DR excitation (Excite). **f**, d-prime, psychometric threshold, and reaction time computed from analyses in e. d-prime and reaction time computed at coherence = 0.2. Activating DR deteriorates performance. **g**, Response bias and lapse computed from the data shown in e. **h**, Average difference sensitivity maps computed from False alarms prior to optogenetic stimulation (Ctrl), FAs preceded by optogenetic stimulation (LED On), and FAs that were not preceded by optogenetic stimulation (LED Off, n=7 mice). Behavioral sensitivity to the cued screen dropped during optogenetic stimulation of DR. Color scale is 0 – 0.4 **i**, Average attentional index for conditions shown in h, also dropped when the DR was stimulated. **j**. % Hits versus stimulus coherence for unstimulated (Control) and Jaws-evoked DR inhibition (Suppress). **k**, d-prime, psychometric threshold, and reaction time computed from analyses in j. Suppressing DR enhances performance. **l**, Response bias and lapse computed from the data shown in j. **m**, Average difference sensitivity maps computed from Ctrl, LED ON, and LED OFF False alarms (n=7 mice). Color scale is 0 – 0.4. **n**, Average attentional index for conditions shown in m. Suppressing DR increases attention to the cued screen. * = P < 0.05, Wilcoxon signed-rank test.

Exciting DR-5HT neurons significantly increased psychometric threshold and reduced d-prime for 3-bar gratings with no effect on reaction time (**Fig. 4e-f, Extended Data Fig. 4a**). These effects arose without change in response bias, lapse rate, and criterion (**Fig. 4g, Extended Data 4c**), indicating that DR excitation reduces sensitivity to gratings rather than affecting impulsiveness, engagement, or strategy. Next, we compared sensitivity difference maps and attention indices across the 3 false alarm conditions. We saw weaker difference maps and significantly reduced attention when the DR was high (**Fig.4h-i**, *Ctrl* vs *LED on*). Moreover, we saw the same weakened difference maps and attention between stimulated and unstimulated FAs during optogenetic sessions. (**Fig.4h-i**, *LED on* vs *LED off*). Thus, elevated DR-5HT activity selectively reduces visual attention to features matching the 3-bar grating and deteriorates behavioral performance.

Suppressing DR-5HT neurons had the opposite effect and significantly increased d-prime and decreased psychometric threshold with no effect on reaction time (**Fig. 4j-k, Extended Data Fig.4b**). Again, these improvements arose without significant changes to response bias, lapse rate and criterion (**Fig. 4l, Extended Data Fig. 4c**), suggesting that low DR activity increased behavioral sensitivity to grating-like features rather than changing impulsivity, engagement, or strategy. Attention indices and sensitivity difference maps were significantly elevated with respect to pre-optogenetic sessions (**Fig. 4m-n**, *Ctrl* vs *LED On*) or with respect to unstimulated FAs (**Fig. 4m-n**, *LED On* vs *LED Off*). No performance or sensitivity changes were seen when we optically stimulated sham control mice (**Extended Data Fig. 4d**). Taking all optogenetic studies together, we conclude that DR-5HT neural activity modulates visual attention and behavioral performance.

### V1 serotonin release dynamics predict visual attention and behavioral performance

Since prior work indicated that attentional selection is implemented by local circuitry in V1^9–11^, we next asked if our DR-5HT attention signal appears in this cortical area. To do this, we delivered AAVs bearing the genetically encoded 5HT sensor GRAB5HT^40^ into V1 and wide-field imaged fluorescent signals while mice performed our task (**Fig. 5a**). Tissue sections through V1 showed broad expression of GRAB5HT across cortical laminae **(Fig.5b)**, and GRAB5HT signals showed similar task-related signals as in our photometry dataset (**Extended Data Fig. 5a**).

**Figure 5.**
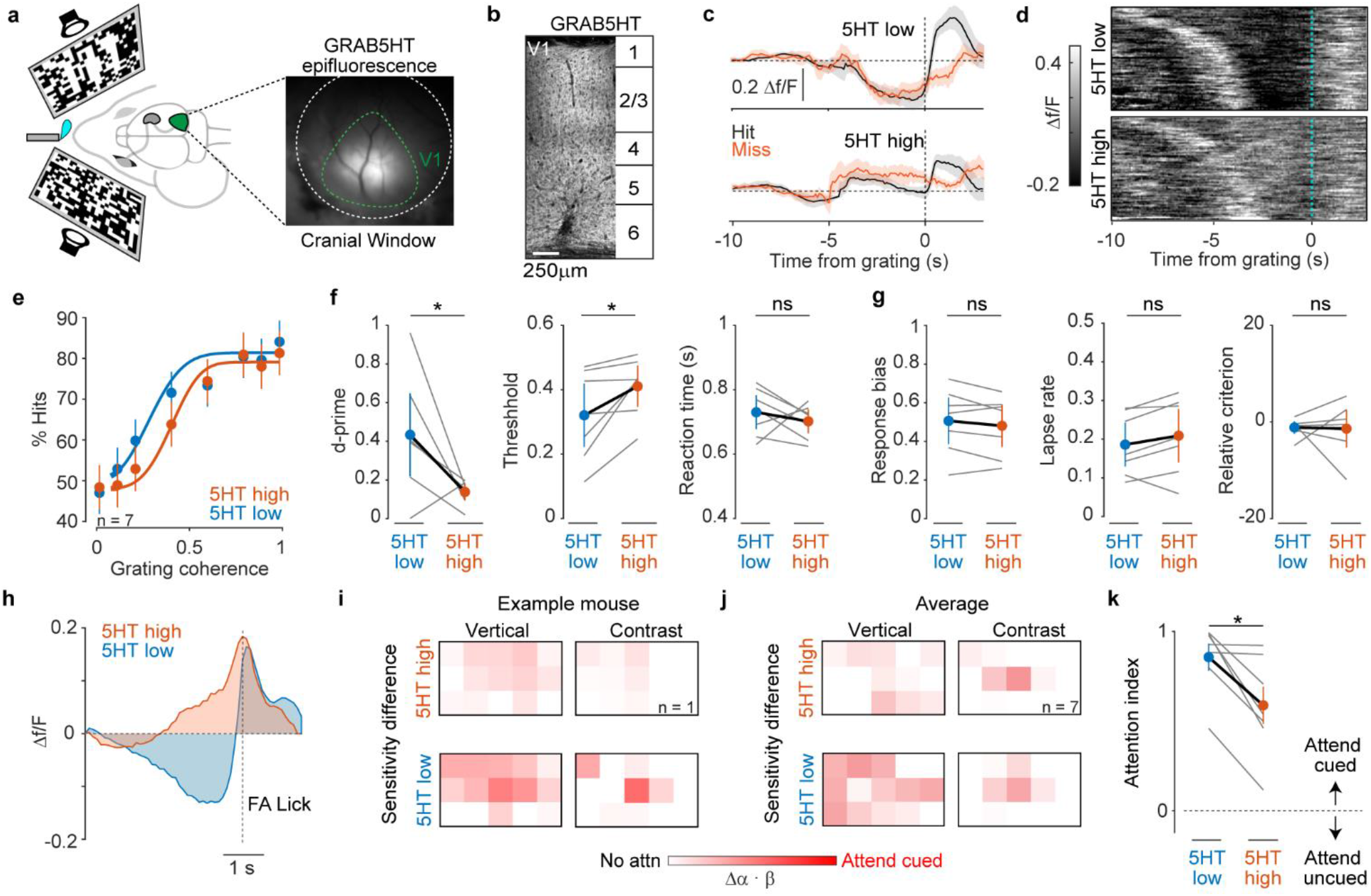
Low 5HT release in V1 predicts attention and detection of the 3-bar grating. **a-b**, Cartoon showing expression of GRAB5HT in V1 while mice detect 3-bar gratings (a), and an example image through a cranial window; V1 border labelled. **b**, V1 tissue section showing GRAB5HT expression throughout cortical depth. **c**, Average GRAB5HT signals for miss and hit trials with low (5HT-Low) and high (5HT-High) DRN. **d**, Raster plots from one mouse showing raw GRAB5HT signals from 5HT-Low and 5HT-High groups. Trials were sorted by the length of the delay period. **e**, Psychometric curves from low GRAB5HT (5HT low) and high GRAB5HT (5HT high) groups. **f**, Average values for d-prime, psychometric threshold, and reaction time computed from data in e (n=7 mice). **g**, Response bias, lapse rate, and criterion computed from data in e. **h**, Average GRAB5HT high and low signals preceding a False-alarm lick. **i-j**, Example (i) and average (j) difference sensitivity map computed from 5HT high and 5HT low FAs. Attention to the cued side increased when 5HT release in V1 was low. Example and average color scales for DR high and DR low are all 0 - 1. **k**, Average attention index from data shown in l (n=7). Behavioral attention increased during low 5HT release. * = P < 0.05, Wilcoxon signed-rank test.

Importantly, we were able to group trials into 5HT low and high trials, with 5HT release signals resembling our DR signals from photometry (**Fig. 5c-d**). Grouping trials according to high or low pre-grating GRAB5HT signal revealed significantly higher hits and d-prime, and lower psychometric thresholds when GRAB5HT signals were low, but no change in reaction time (**Fig. 5f**). These effects arose without significant changes to the response bias, lapse rate, and criterion, suggesting low 5HT release is associated with better stimulus detection rather than differences in impulsivity, engagement, or strategy (**Fig. 5g**). Taken together these results show that low 5HT in V1 predicts better performance.

We next estimated behavioral attention from FA licks preceded by either high or low GRAB5HT signals (**Fig. 5h**). Sensitivity difference maps were biased towards the cued side when GRAB5HT was low versus when it was high (**Fig. 5i-j, Extended Data Fig. 5b**). Moreover, the average attention index was ∼25% greater during GRAB5HT low as compared to GRAB5HT high (**Fig. 5k**). These results indicate that V1 5HT release predicts the strength of visual attention and behavioral performance. Taken together with our photometry and optogenetic results, we conclude that DR-5HT is a novel visual attention signal.

## Discussion

Visual attention is viewed as a spotlight that enhances the most behaviorally relevant stimuli^1,2^. While progress has been made towards the cortical contributions to this spotlight^9–11^, the long-suspected contributions^13,18,41^ of subcortical neuromodulatory structures remain elusive. Here, we present evidence that DR-derived 5HT is a significant source of attentional regulation in mice. Using fiber photometry and optogenetics, we show enhanced visual attention and detection when DR-5HT neuron activity and 5HT release in V1 is low. When DR-5HT neuron activity and V1 5HT release are high, mice showed reduced attention and performance. This effect of 5HT was specific to stimulus detection and did not alter measures of impulsivity, engagement, or strategy. Taken together, these results define DR-5HT as a new member of the brain’s attentional network.

### How does DR-5HT modify attentional networks?

Our data consistently showed an inverted relationship between 5HT and attention that agrees with prior work on the suppressive effect of 5HT in V1^23–26^. In mice, 5HT receptors expressed by V1 pyramidal cells and their LGN inputs (depressive 5HTR1 types^29,42–44^) along with those expressed by V1 interneurons (stimulatory 5HT2a^24,27,45^) predict a net inhibitory effect for of 5HT. This matches older *in vivo* studies in non-human primates showing that 5HT iontophoresis suppressed response gain of V1 neurons^25,26^. Our results also agree with newer studies in mice showing that 5HT attenuates visual responses on individual V1 neurons^23^ and divisively scales V1 population responses to stimuli^24,27^. Such scaling would represent a change in gain which is believed to be central to attentional computations. Taking these studies together with our results predicts that a drop in 5HT would multiplicatively scale V1 neurons’ tuning to visual stimuli. We previously saw such scaling of behavioral tuning curves in this same task^36^, simultaneously measuring tuning curves and 5HT release from individual V1 neurons could shed more light on this issue.

### DR 5HT-signals regulate the strength of attention

Our study explicitly modeled attention as a gain term (α) applied to a map of sensitivities to visual locations and features (β, see **Fig. 1d**). DR-5HT high and low activity produced significant changes in the strength of attention, but did not change the locations and features mice monitored to perform the task (**Figs. 1, 3-5**). It is as if 5HT changed the strength of the attentional spotlight but did not change where and what it shines upon. Why would the brain employ 5HT to regulate attention in this way?

One reason could relate to the brain’s use of visual cues to guide attention. The environment often contains many potential cues, and each can vary in their predictiveness over time. Thus, a cue engages attentional circuits only if the brain deems them valid enough to predict future events. Could our DR-5HT signal reflect cue validity and thus modulate attention gain rather than modulating sensitivity to visual locations and features? Older work on another neuromodulator, acetylcholine (ACh), supports this view^12,16,46–49^. In these studies, ACh encoded cue validity by tracking another parameter, expected uncertainty^50^. ACh is high when the outcome (or uncertainty therin) is predicted by a cue; ACh was low when unexpected shifts in context stole predictiveness from the cue^46,50,51^. Interestingly, recent work indicates that tonic DR-5HT firing is low when expected uncertainty is high^52,53^. These results suggest that cue validity could be encoded in DR-5HT activity. When low, the cue is valid, and attention is strong.

In our task, however, the audiovisual cue was always valid. So why did attention and DR-5HT fluctuate? One possibility is that grating-like patterns randomly produced by checkerboard noise on the uncued side and/or block switches stole predictiveness from the cue. Comparing expected and unexpected uncertainty over time, as captured by reinforcement learning models^50,52,53^ could shed more light on this issue.

### Neuromodulatory control of attention strength

William James famously wrote “Everyone knows what attention is”^54^. Nearly 75 years later, we still do not fully understand attention in biological terms. Studies in non-human primates, indicate that frontal cortical feedback excites V1 in ways that mimic the effects of attention^9–11,55–61^. Newer studies in rodents have partially identified V1 circuitry that uses such feedback to enhance V1 neural responses^8–11,62–65^. However, a separate body of work suggests that subcortical neuromodulatory areas are a major uncharacterized source of attentional modulation^13,18,41^. The data presented here provides strong support for this view. Interestingly, frontal cortical areas implicated in attentional feedback to V1 represent the strongest inputs to the DR^19–21,37^. Retrograde tracing from DR and V1 could reveal whether such frontal cortical ‘attention’ neurons represent exclusive or overlapping populations.

A small set of studies suggest that 5HT influences human attention^66–69^. A study which depleted 5HT and showed improved attention^66^ is consistent with the data we have shown here. Screening the many drugs that target 5HT signaling in our task could reveal new routes to improve the weakened attention that characterizes several disorders including attention-deficit hyperactivity disorder and autism.

## Methods

### Animals

Male and female Cdh13-CreER (Cdh13^Ce^), ePet1-Cre (ePet1), Ai95 and Ai27 mice were used in this study. Rosa26-LSL-ChR2-tdTomato (AID27, RRID:IMSR_JAX:012567) and Rosa26-CAG-GCaMP6f (RRID: IMSR_JAX:028865) mice were purchased from the Jackson Laboratory and crossed with ePet1-Cre mice or Cdh13-CreER for recording and optogenetic experiments, respectively.

For photometry recordings we treated three Cdh13^Ce/+^-G6f^+/+^ mice with tamoxifen to drive expression of GCaMP6f. The remaining 6 animals expressed GCaMP6f under the control of *Pet1* promotor driven Cre expression. Optogenetic excitation studies used four Cdh13^Ce^-Ai27D and four ePet1-Ai27D animals. Optogenetic suppression studies were performed on 7 ePet1 animals that received 1ul of AAV-Jaws (*see Injections*). Imaging of 5-HT release in visual cortex was performed on 7 C57BL/6J wild-type mice. All surgical and experimental procedures were in accordance with the rules and regulations established by the Canadian Council on Animal Care, and protocols were approved by the Animal Care Committee at McGill University.

### Tamoxifen

Tamoxifen (TMX) was dissolved in anhydrous ethanol at 200 mg/mL, diluted in sunflower oil to 10 mg/mL, sonicated at 40ºC until dissolved, and stored at –20°C as previously described^70,71^. Prior to injection, TMX aliquots were heated to 37°C and a single injection was delivered intraperitoneally at ∼1 g/50 g body weight to Cdh13-CreER ∼P30 mice.

### Behavioral arena

We used the same behavioral arena as described previously^36^. Briefly, the behavioral apparatus consisted of a custom built soundproofed dark box that displayed visual stimuli via two 60 Hz LCD displayed at a 32-degree angle from the mouse’s midline. The mouse was head fixed in the center of the apparatus on a platform and a lick spout positioned to administer liquid reward. Licking was capacitively sensed and digitized using custom electronics. Liquid reward delivery was controlled with a solenoid pinch valve driven by custom electronics. The behavioral task and presentation of visual stimuli was controlled using MATLAB’s psychophysics toolbox (Psychtoolbox3), and custom data acquisition software.

### Behavioral task design

We trained mice in our task as described previously^36^. Briefly, mice performed a spatially cued detection task and were trained to lick in response to a 3-bar grating embedded within a noisy checkerboard background. The grating coherence was parametrically controlled by altering the number of noisy checkers that the 3-bar grating contained (**Fig.1b**). At a grating coherence of 1, the 3 bars contain no noisy checkers, although checkers surrounding the 3 bars are always noisy. At a grating coherence of 0, all checkers are pure noise. While the 3 vertical white bars in the grating increased vertical and local contrast energy, the total number of white checkers in the checkerboard stimulus was always 50%.

Trials began with an auditory cue and a static random checkerboard. Mice would lick to start the dynamic checkerboard noise. The coherent grating then occurred on the cued screen at a random time, chosen between 3-12 seconds. For each individual trial, the wait time before the coherent grating appeared was randomly chosen to produce a constant probability of appearing^36^ (i.e., a flat hazard function, **Fig.1d**).

A lick that occurred within 0.1 - 1.5 seconds following the 3-bar grating was labeled as a hit and led to rewards. The absence of licks during this window were misses and were not rewarded. If mice licked before the onset of the 3-bar grating (false alarm), the trial was restarted. Restarts caused the audiovisual cue to reappear for 0.5s, followed by a return of dynamic checkerboard noise.

### Training schedule

Reverse-cycled (12hr/12hr inverted light-dark) mice were trained as previously described^36^. Briefly, water restricted mice were habituated to behavioral arenas for 2 days prior to behavioral shaping. Our shaping protocol is as follows: (1) 1 day to associate licking with liquid reward; (2) 4 days to associate strong 3-bar gratings (coherence> 0.8) with reward; and (3) 7-10 days to lengthen maximum wait times to 12 seconds and decrease grating coherences. By session 15, mice produced reliable psychometric curves. A typical training session lasted 1.5-2 hrs. Mice were never rewarded for false-alarm licks, or licks in response to a zero-coherence grating.

### Behavioral analysis of grating detection

For photometry experiments, we pooled trials from later sessions (>session 15) and divided them into DR-low and DR-high categories before computing psychometric measures of performance. For optogenetic studies, control datasets consisted of 10-15 sessions prior to LED stimulation; experimental datasets consisted of all trials during LED stimulation.

Psychometric functions for hits as a function of grating coherence were described by a Weibull function, which provided four parameters: mean, threshold, response bias (lower asymptote), and lapse rate (higher asymptote). Here, threshold represents the grating coherence that is detected at 50% rate of the total response range. Signal detection theory was used to compute the discriminability index d-prime = *Z*(*PH*) ™ *Z*(*PH*0) and criterion = ™ 0.5 *X* (*Z*(*PH*) *+ Z*(*PH*0)) / d-prime, where *PH* is the probability of a hit at a single grating coherence. *PH*0 is the probability of a zero-coherence hit, and Z is the inverse of the standard normal cumulative distribution function. Reaction times were described by a standard exponential function: *ae*^™*b* (*coherence*)^ *+ c*, with amplitude (*a*), baseline (*c*), and slope (*b*).

### Quantification of behavioral sensitivity and attention

Our previously published logistic model revealed that attention multiplicatively scaled mouse behavioral sensitivity for grating-like features on the cued versus the uncued side^36^. Our current model simplifies this approach to capture this attentional scaling in a single parameter. Briefly, we used false-alarm licks preceded by at least 1-3 seconds of lick-free random checkerboard noise from the final 20 sessions of mice. This removed FA lick “bouts” and FA licks occurring before the reaction time window but after grating presentation (FA outcomes).

Features were computed as follows: Vertical stimulus energy was computed at the same spatial frequency as the target stimulus (0.077 cycles per degree) by convolving quadrature phased Gabor filters (26×26 checkers, S.D = 6 checkers or 3 deg of visual angle) with each stimulus frame. Contrast energy was computed by convolving each stimulus frame with a Gaussian filter (S.D. = 4 checkers, 2 deg).

Resulting energy maps for the two features were then down sampled in time from 30 Hz to 10 Hz and spatially down sampled from 17×26 checkers to 3×5 checkers to ease computational load. Energies for both features were z-score normalized.

FA licks were modeled using the logistic function:

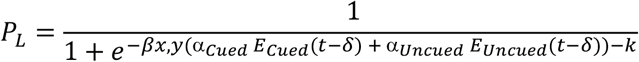

In this equation, the probability of a FA lick (*P*_*L*_) is a function of z-scored stimulus energy, characterized as vertical or local contrast. In this model, δ is the time it takes for stimulus energy to produce a FA lick, chosen as a hyperparameter based on each individual mouse’s psychometric reaction time distribution; we previously showed individualized reaction times for mice^36^. We optimized 3 unknown model parameters: *βx, y* –the sensitivity of FA licks to the stimulus energy on both the cued or uncued screen for each individual checker at location (*x, y*); *θ* –the attentional gain parameter (constrained to [0, π/2]) which determines the weighting of cued and uncued stimulus energy (*E*_*Cued*_ and *E*_*Uncued*_,), and *k* –the stimulus energy corresponding to *P*_*L*_ = 0.5. The weighting of cued and uncued energies were captured by the relationship:

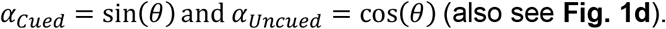

The model parameter returns can be interpreted as follows: β is a spatial distribution of behavioral sensitivity applied to both the cued and uncued screens, and *θ* is an attention gain that scales energies from cued side if *θ* > π/4, such that α_*Cued*_ >α_*Uncued*_. Both β and *θ* were optimized to maximize the model predictive power over FA behavior (**Extended Data Figure 1a**). Attentional gain *θ* = *pi*/4 means no attention to cued features over uncued features, and *θ* = *pi*/2 is 100% attention to cued features over uncued features.

We treated the model as an optimization problem for a constrained nonlinear multivariable function, implemented using MATLAB’s *fmincon* with the negative-log-likelihood cost function

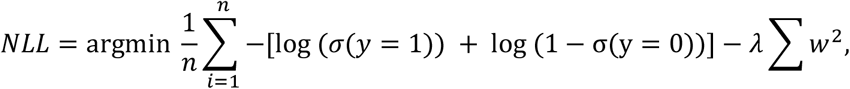

where λ is the regularization factor for standard L2 regularization, *w* are the parameters β_x,y_ , *y* are the *n* observations (lick or no lick), and σ the standard logistic function. Each model was 5-fold cross-validated to optimize hyperparameters without bias. Model predictive power was measured using the area under the receiver-operator-characteristic.

### Fiber optic implants

For photometry and optogenetic studies, mice were anesthetized (isofluorane, 4% induction, 1-2% maintenance) mounted on a stereotaxic frame and eye ointment was applied to prevent corneal drying. Following lidocaine application in subcutaneous carprofen injections, a circular incision was made, and the underlying fascia cleaned to dry the skull. Bregma and lambda were then levelled to within 0.1 mm of one another. A 0.1 mm craniotomy was performed above the DR, and the fiber implant (400 μm diameter fiber, ThorLabs) was slowly inserted using a stereotactic arm (4.7 mm posterior from bregma, on the midline and 3.1 mm below the pia). Metabond cement was applied around the implant location and a headplate was cemented additionally to the skull. Any remaining exposed skull was sealed using cement. Following surgery, the animal was removed from the stereotaxic frame and placed on a heat pad for recovery, and analgesia was administered subcutaneously following surgery for 3 days.

### Cranial window

For GRAB5HT imaging in V1, mice were anesthetized and mounted on a stereotaxic frame as described above but received a 5 mm diameter craniotomy over the left V1 followed by a circular glass coverslip (4 mm diameter) implant centered 1 mm rostral from lambda and 2.5 mm latera from the midline. Cranial windows were fixed using instant glue and Metabond at a 15^°^ angle to the mediolateral axis and then received a metal headplate implant that surrounded the craniotomy.

### Injections

AAVs were intracranially injected using previously described methods^72,73^. Mice used in the optogenetic suppression experiments received a 1 ul injection of AAV1-CAG-FLEX-rc [Jaws-KGC-GFP-ER2] in the DR (4.7 mm posterior from bregma, on the midline and 3.3 mm below the pia). Animals for 5-HT imaging received three 500 nl infusions of AAV9-hsyn-GRAB_5-HT1.0 in the left visual cortex (centered around 1 mm rostral from lambda, 2.5 mm latera from the midline at a 30^°^ injection angle and 0.2 mm below the pia). A microsyringe pump (UMP3-4, World Precision Instrument, Sarasota, FL) was used to infuse virus at 5 nL/s.

### Viruses

AAV9-hsyn-GRAB_5-HT1.0 (Cat# 140552-AAV9) and AAV1-CAG-FLEX-rc [Jaws-KGC-GFP-ER2] (Cat# 84445-AAV1) were purchased from Addgene and used for intracranial injections.

### Photometry

A 1-site custom-built fiber photometry system was used to assess changes in GCaMP6f fluorescence from DR neurons. Calcium insensitive fluorescence was measured with 405 nm light oscillating at a carrier frequency of 450 Hz. Calcium sensitive fluorescence was measured with 472 nm light oscillating at 205 Hz. An optical fiber (ThorLabs) was used to both deliver and collect light from the brain. Light emitted in the 500-550 nm band was measured using a femtowatt photoreceiver and digitized 200Hz with an RX8 signal processor (Tucker Davis Technologies). The recorded signal was demodulated according to the two carrier frequencies and low pass filtered at 20 Hz. Calcium signals were discretized into individual sessions and calcium insensitive recordings were fit to the calcium-sensitive signal using least-squares regression using a 5-minute moving window. The fit calcium insensitive signal was subtracted from the total signal. The resulting signal was normalized to the mean of each session via z-scoring to yield a final measure of ΔF/F.

### Wide-field imaging

We used a custom-built epifluorescent microscope to image through our cranial windows with a 10 Hz sampling rate at 512×512 pixel resolution. Since windows were mounted at a 15-degree angle with respect to the mouses medio-lateral head axis, we mounted the microscope at the same angle such that mouse heads remained leveled with respect to the visual stimuli. GRAB5HT fluorescence was measured with a 472 nm excitation LED and LabVIEW custom code. Session signals were z-scored pixel-wise to yield a final measure of ΔF/F. Signals in **Fig. 5** are pooled averages of the brightest 10% of pixels in the image.

### Photometry signal processing

To explore how DR activity on short timescales affected animal performance in our detection task, we first high pass filtered our ΔF/F session signals recorded via fiber photometry or widefield imaging with a cutoff frequency of 1/40 Hz. Since only hit trials contained reward events that accompanied large positive DR reward signals (see **Fig. 2f**), a filter may artificially introduce baseline shifts preceding the grating presentation on hit trials, but not on miss trials. To eliminate these potential baseline artifacts preceding grating presentation, we removed reward signals from our data before filtering. Doing so ensures that hits and misses were treated the same. We then computed the average baseline activity in a window preceding grating onset for each trial; this time window was half the length of the unique delay period for each trial. Trials below the median baseline activity of all trials were categorized as DR low, trials above the median were categorized as DR high. False alarms were similarly grouped into DR low and DR high based on the mean activity of 1.5 seconds preceding the false alarm.

For long time-scale analysis (**Extended Data Figure 3**), session signals were low pass filtered using a 5 Hz cutoff frequency. Trials in the session were then identified as DR low if most of the trial signal (80% of data samples) remained below the median of the session signal. Trials above the session signal were labeled as DR high.

### Optogenetics

We developed custom code on an Arduino Due to drive a high-powered LED (Thorlabs) that was attached to the mouse’s implant through an optical fiber (400μm diameter, Thorlabs). Our custom behavioral code was adjusted to have full control of the LED stimulation time. For experiments involving mice expressing ChR2, we drove a 470 nm LED at 20 Hz (150 ms pulse duration, 17.2 mW). Mice carrying AAV1-Jaws, received a constant 595 nm LED input (8.7 mW) for the duration as suggested previously^74^. Mice were trained fully on the visual detection task (30-40 sessions), and then received LED stimulation on every trial for at least 15 more sessions. LED stimulation began 2 seconds before the grating stimulus was scheduled for a duration of 3 seconds. False-alarm licks could interrupt LED stimulation and led to trial restarts as described above.

### Histology

Mice were euthanized by isoflurane overdose and perfused intracardially with phosphate buffered saline (PBS), followed by 4% paraformaldehyde (PFA) dissolved in PBS chilled to 4 degrees. Brains were removed and postfixed in 4% PFA overnight. Next, brains were embedded in 2.5% agarose (Sigma), mounted on a tissue slicer (Compresstome), and 150 um thick coronal brain sections were sliced and stored in PBS until immunostaining. Brain sections were incubated with blocking buffer (10% normal donkey serum, 0.4% Triton X-100 in PBS for 1-2 hours), then incubated for 7 days at 4°C with primary antibodies, and with secondary antibodies overnight at 4°C. Primary antibodies were used as follows: rabbit anti-TPH2 (1:1000, Novus; 1:500, Synaptic Systems); chicken anti-GFP (1:1000, Abcam). Secondary antibodies were conjugated to Alexa Fluor 488 (Invitrogen), or Cy3 (Millipore). Sections were mounted onto glass slides using Vectashield (Vector Lab) or Fluoromount (Millipore) mounting medium.

Immunostained images were acquired from a Zeiss confocal Microscope, using 405 nm, 488 nm, 559 nm, and 635 nm lasers. ImageJ (NIH) software was used to analyze confocal stacks and generate maximum intensity projections to one image.

## Contributions

JL, AK, and EC conceived of this study and aided with instrumentation, analysis, experiments, writing, and making figures. JB provided the photometry system and gave experimental advice. JL and KC devised the model with guidance from EC and AKr. KY, JL, and XM helped with behavioral training.

## Acknowledgements

We thank A. Rangel Olguin, S. Shariff, N. Brake, and F. Jalondoni for helpful discussions and contributions.

## Extended Data Figures

**Extended Data Figure 1.**
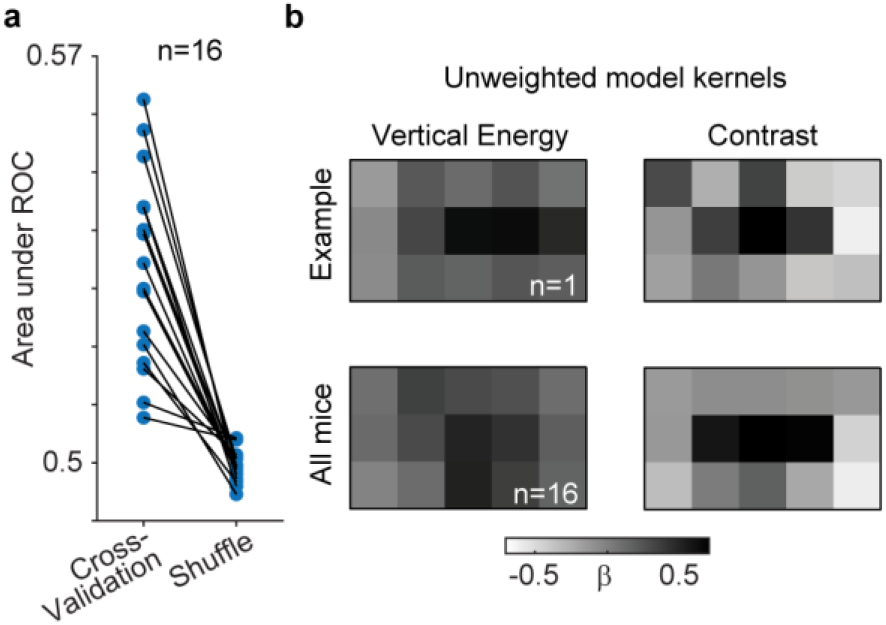
Attention Model Spatial filters and Validation. **a**. Cross-validated area under the receiver operator curve (ROC) for 16 mice compared to model fits in which checkerboard sequences were shuffled relative to false alarm licks. **b**. Model sensitivity maps and for vertical (βv) and contrast (βc) energy obtained from an example mouse (top) and average model sensitivity maps from 16 mice.

**Extended Data Figure 2.**
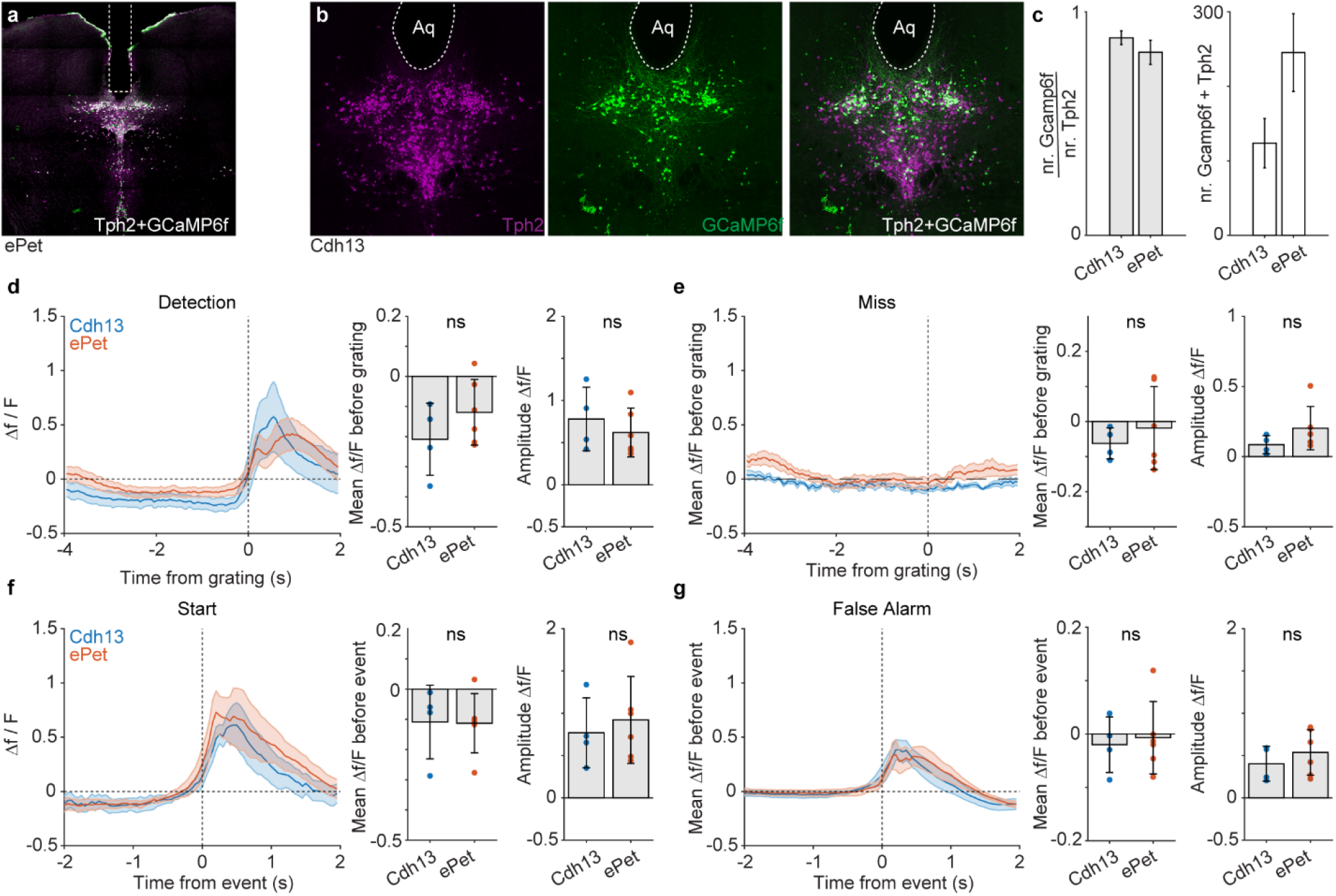
Comparison of GCaMP6f signals from Cdh13-CreER and epet1-Cre mice, **a**. Example coronal section through the DR of an epet1-Cre x Cre-dependent GCaMP6f (Ai95) mouse stained with antibodies against tryptophan hydroxylase 2 (Tph2) and GFP. **b**. Example coronal section taken from a Cdh13-CreER x Ai95 mouse stained for Tph2 (left) and GFP (middle). Right image shows the merge. **c**. Fraction (left) and absolute number of Tph2+ neurons that are also GCaMP6f+ in epet1-Cre x Ai95 and Cdh13-CreER x Ai95 mice. Both lines label similar numbers of Tph2+ neurons with GCaMP6f but epet1-Cre produces more complete labelling. **d-g**. Average GCaMP6f photometry signals from epet1-Cre or Cdh13-CreER mice aligned to detection (d), miss (e), initiation (f), and False Alarms (g). Bar graphs at right quantify the mean Δf/f prior to the indicated event and the mean post-event amplitude.

**Extended Data Figure 3.**
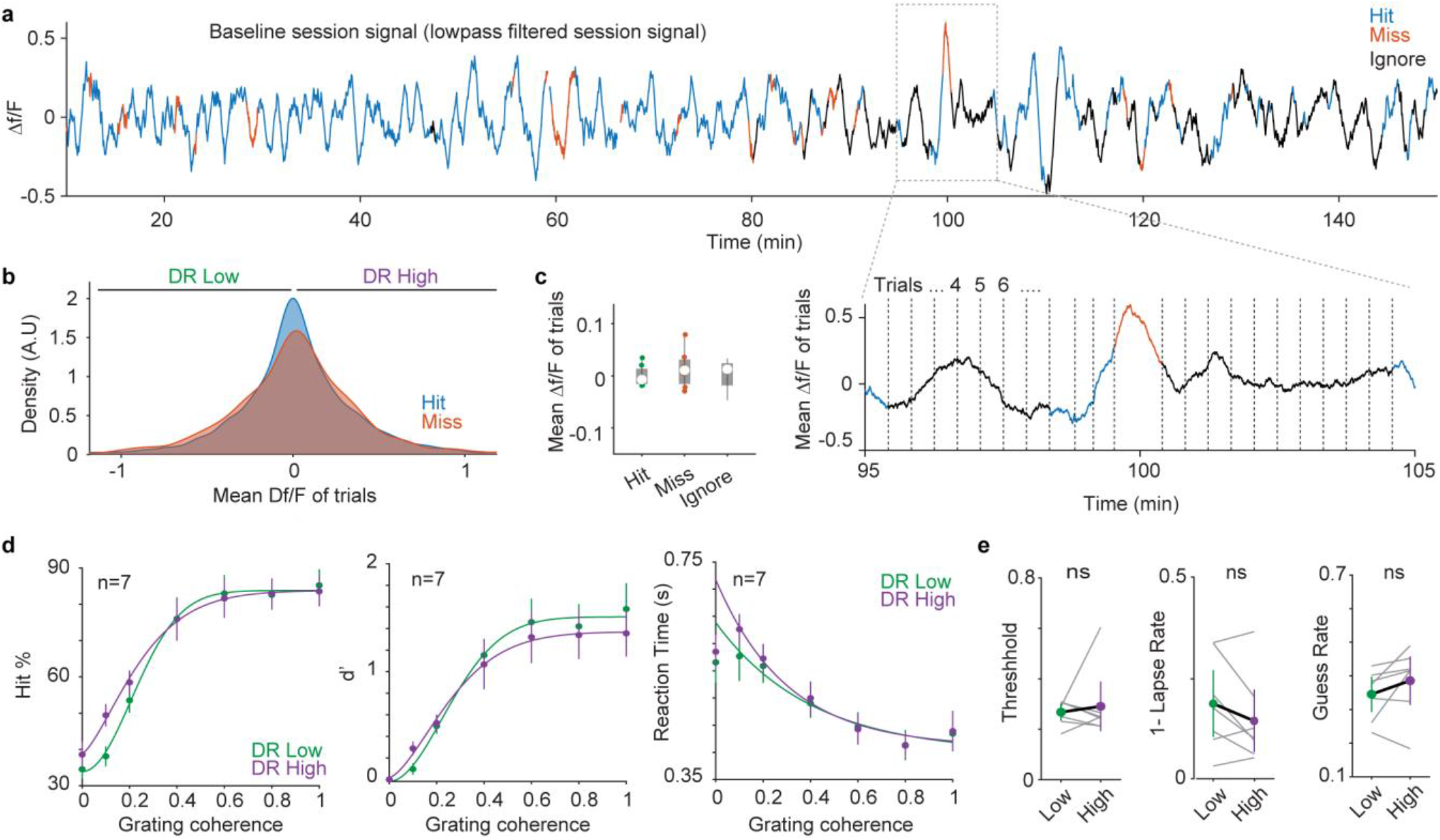
Long-timescale DR dynamics and behavioral performance, **a**. Example trace showing change in DR baseline activity across ∼150 minutes. Trace is colored according to hit, miss and ignore trials. Inset magnifies the boxed region. **b**. Histogram of baseline Δf/f during hits and misses taken from session signals like shown in a. **c**. Average Δf/f for Hit, Miss, and Ignore trials. **d**. Psychometric curves of Hit rate, d-prime, and reaction time versus grating coherence **e**. Average psychometric threshold (left), lapse rate (middle) and FA rate (right) computed from data like shown in d. n> 1500 baseline segments from 7 mice.

**Extended Data Figure 4.**
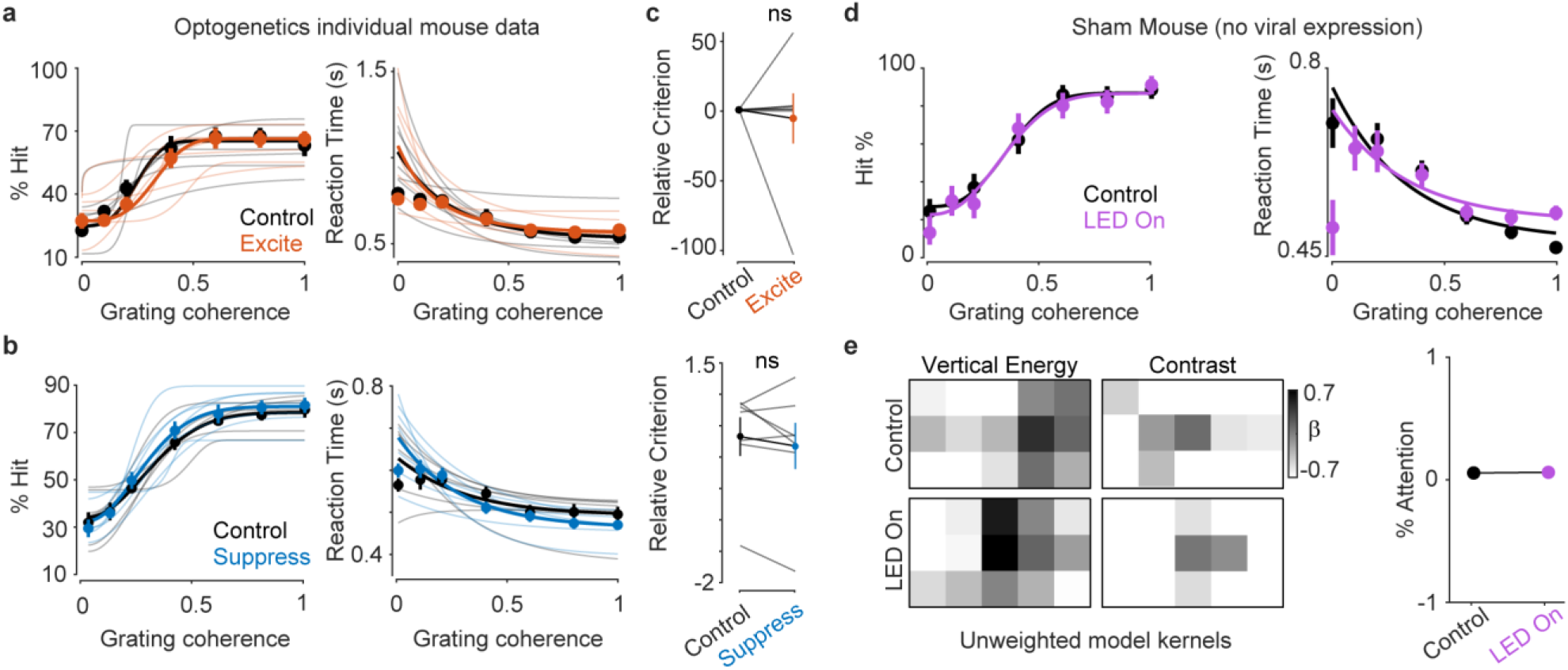
Mice for optogenetic experiments were trained to similar performance level as controls, **a-b**. % hits (left) and reaction time (right) versus grating coherence computed from control trials or trials with optogenetic excitation (a) and suppression (b). **c**. Relative criterion computed from experiments like shown in a-b and in Figure 4e and j. **d**. % hits (left) and reaction time (right) versus grating coherence computed from control trials or trials with optical excitation in sham mice. **e**. Model sensitivity maps (left) and attention (right) for sham experiments shown in d.

**Extended Data Figure 5.**
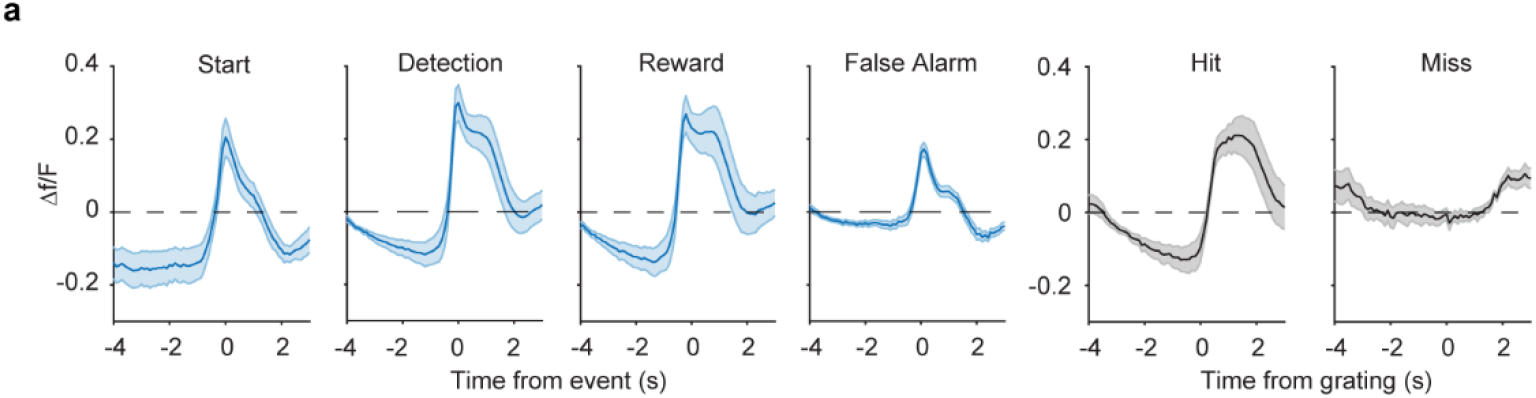
GRAB5HT signals from V1 aligned to task events and outcomes, **a**. GRAB5HT signals aligned to start, reward, and false alarm licks (task events), and to hits and misses (task outcomes).

